# Gene-family-dependent thermodynamic effects of oncogenic mutations on nucleosome-DNA binding stability: a comparative molecular dynamics study across 22 cancer hotspots

**DOI:** 10.64898/2026.09.28.754892

**Authors:** Shlomo Freudenberger, Evan Boczko, Sanjay Goel, Radhashree Maitra

## Abstract

Oncogenic mutations are known to occur at non-random rates and specific hotspots, yet the structural and thermodynamic factors underlying these hotspots remain poorly understood. This study presents molecular dynamics simulations of 22 cancer hotspots across 10 oncogene families, simulated as histone-DNA complexes. Across 18 of 22 mutations, thermodynamic effects were predominantly localized to within 6 Å of the DNA-histone interface (mean capture 104%), with van der Waals interactions driving the effect at the thermodynamic extremes, indicating that oncogenic mutations alter nucleosome binding through precise local contact changes rather than global structural rearrangements. Further, analysis revealed a gene-family correlated pattern in thermodynamic stability. RAS family mutations showed a consistent trend toward nucleosome stabilization (mean ΔΔG = −4.82 kcal mol^-1^), while kinase domain mutations trended toward destabilization (mean ΔΔG = +40.12 kcal mol^-1^), a difference reaching statistical significance in this exploratory analysis (Mann-Whitney U, *p* = 0.008). This interface-specific mechanism, combined with the gene-family-correlated thermodynamic pattern, provides a biophysical framework for understanding nucleosome-level contributions to cancer hotspot mutation biology.

## Introduction

### 1.1 Genomic hotspots and the chromatin context

DNA mutations are a hallmark of cancer progression [1]. It has been established that cancer mutations are not randomly distributed throughout the human genome, rather are statistically found at higher rates at specific points in the sequence, leading to known hotspots [2].

Specifically, there are codons which demonstrate heavy recurrence of mutations, including distinct amino acid substitutions at the same codon in different cancers [3]. Studies have elaborated on the selective advantage that specific mutations provide to the cancer cells [4]. These mutations are positively selected for during the evolution of the cancer.

Much research has explored short DNA sequences that are prone to mutate through distinct biochemical processes [5–8]. But not only are DNA mutations implicated in tumor progression, but epigenetic modifications controlling gene expression are also relevant. Feinberg and Vogelstein demonstrated that human carcinomas exhibited widespread loss of 5-methylcytosine at CpG sites compared to adjacent normal tissues [9]. Furthermore, methylation patterns can lead to regions of hypermutation, referred to as kataegis – Greek for shower or thunderstorm [10].

Another factor in cancer progression is the cells various internal DNA repair mechanisms. While these mechanisms are not isolated to the study of cancer, they most certainly play a role in the prevention of cancer progression [11]. Interestingly, DNA accessibility within the chromatin structure may be implicated in tumorigenesis due to inhibition repair mechanisms [12].

While it has been documented that sequence-based studies can provide indications of cancer hotspots, there remains a clear gap in the three-dimensional structure and thermodynamic context of these mutations within the chromatin. Mutations do not occur in isolation. They occur within the context of DNA wrapped around histone proteins, where mutations can alter the stability of DNA-protein interactions. This may influence the likelihood of a mutation occurring and/or its likelihood of being retained through selection.

### 1.2 Nucleosome structure and DNA-histone thermodynamics

Nucleosomes are the most basic structure of condensed DNA found in the cell. A family of histone proteins, including the core proteins (H2A, H2B, H3, H4) and a linker protein (H1), form a protein complex. Note that the linker histone H1 is not present in the nucleosome core particle used in this study (PDB: 2CV5); all simulations model the histone octamer without linker histone. The nucleosome consists of approximately 146 base pairs wrapped around an octamer. Histone proteins are rich in basic, positively charged lysine and arginine residues which form strong electrostatic interactions with the negative charged phosphate backbone of DNA. DNA geometric shapes, such as minor groove width, can also modulate the strength of the electrostatic interactions [13].

The location of a sequence within a nucleosome is not a random process. Rather, DNA sequences have higher thermodynamic stability within different specific regions of a nucleosome [14]. This provides evidence that DNA position along a nucleosome is not considered a random process, rather DNA will minimize free energy in the most stable thermodynamic position.

These factors, along with many others, provide a continuum of thermodynamic stability or instability present in DNA and histone interactions. Stability is not binary, existing as either stable or unstable, rather there is a spectrum of increasing thermodynamic stability. And even modest effects can have profound impacts on thermodynamic stability. This is also particularly relevant to cancer, as evidenced by Pich et al [15], where periodic patterns are found in cancer mutations.

### 1.3 MM-GBSA as a tool for comparative nucleosome energetics

GROMACS [16,17] is an open-source molecular dynamics (MD) simulation software. It simulates the application of Newtonian physics of motion for biological systems. It is well equipped to handle complex interactions between macromolecules, such as DNA and proteins, as present in this study.

MM-GBSA [18] is an established method to calculate free energy, used in conjunction with GROMACS. It takes conformational snapshots extracted from MD trajectories of binding partners and calculates the binding free energy (Δ*G*_bind_) by removing solvents and then computing the binding. It combines molecular mechanics gas-phase energies (Δ*E*_MM_) with implicit continuum solvent electrostatics (Δ*G*_GB_) and non-polar solvation terms (Δ*G*_SA_) to yield a final binding free energy value. While the absolute values of binding energy express large uncertainties due to the challenge of computing conformational entropy, it is well-suited for comparative screening studies, providing relative ranks which are significant. Since the limitations are systemically consistent it is valid to compare relative levels. Binding energies have previously been validated in analyzing DNA-histone interactions [19–21], but have not been used to explore the relative stability found in cancer mutations in comparison to the wild type sequence. That is the goal of the present study.

### 1.4 The present study and its aims

This study utilized GROMACS to run 200 ns MD simulations of 22 paired mutation sets. Each set consisted of the wild type DNA sequence wrapped around the histone octamer. The mutant version was the same as its respective wild type, except for the substituted point mutation to the DNA sequence. MM-GBSA binding energies were calculated and were used to compare the wild type and mutant of each respective simulation. The ΔΔG landscape was investigated across 10 oncogene families.

Rather than testing a specific directional hypothesis, this study takes an exploratory approach. Are there systemic patterns in nucleosome binding thermodynamics that emerge across a panel of hotspot mutations from diverse gene families? The results reveal a gene-family correlated pattern in thermodynamic direction, suggesting that the biophysical consequences of oncogenic mutations at the chromatin level may be impacted by the class of protein.

## Methods

### 2.1 Structural preparation

For the MD simulations investigating the DNA-Histone interactions, a DNA-nucleosome complex (Supplementary Figure S1) was obtained from the Protein Data Bank (PDB ID: 2CV5). In this complex, the DNA is wrapped around the histones octamer that constitute a nucleosome [22]. This is the appropriate structure because it is the human nucleosome core particle at high resolution with all eight histones.

COSMIC was used to search for an array of “tier one” point mutations to be used in the study [23]. For each of the 22 oncogenic mutations selected, the corresponding wild type sequence was obtained from the NCBI gene database [24]. NuPoP [25], an R-based duration Hidden Markov Model (dHMM) that estimates base pair positions on nucleosomes, was used to predict the location where the mutation is located within the 146 base pairs that wraps around the nucleosome octamer. NuPoP has been validated against MNase-seq nucleosome positioning data. This program is based on research that suggests that the raw DNA sequence itself has a large part in determining the position of the histone proteins along the DNA strand. Once the position on the nucleosome of each selected gene was determined the sequence was precisely extracted from the NCBI database. For example, if a mutation localized to position 100 within the 146-bp nucleosomal DNA, a window spanning 99 base pairs upstream and 46 base pairs downstream was extracted.

The original PDB obtained had 146-base-pair palindromic fragment engineered from human *α*-satellite DNA. Using ChimeraX [26], the predicted base pairs of each gene replaced the native DNA in PDB 2CV5 in the correct position based on NuPoP results. Although the model downloaded from the PDB database did not originally contain the intended gene, the ChimeraX software allows the user to insert any given DNA sequence and replace the default sequence of the PDB model. Thus, a model with the wild-type gene and a model with the mutated gene could be generated and compared in simulations. In total 44 nucleosome core particle models were created (22 wild type and 22 mutant pairs) for subsequent MD simulations.

### 2.2 Molecular dynamics simulation protocol

A total of 44 simulations were run using GROMACS software with identical simulation parameters. Each simulation lasted 200 ns. A simulation length of 200 ns per system was selected based on prior nucleosome MD studies demonstrating convergence of binding energetics within this timeframe [27–29]. Convergence was confirmed for all 44 simulations by RMSD plateau analysis (Section 2.3). A cubic simulation box with 1.0 nm minimum solute-to-edge distance was used, sufficient to prevent self-interaction across periodic boundaries for the nucleosome complex. It was prepared for molecular dynamics simulations using the AMBER14SB/parmbsc1 combined force field, which provides protein parameters from AMBER14SB and DNA backbone parameters from parmbsc1 [30, 31]. The water model TIP3P was used for solvation [32], neutralized with Na/Cl ions to physiological concentration (150 mM). All 17 histidine residues across the eight histone chains were assigned the HIE tautomer (epsilon-protonated neutral form) based on PROPKA pKa predictions at pH 7.4 [33], consistent with the predominantly neutral protonation state expected at physiological pH (pKa values ranged from 3.64 to 7.08). Energy minimization was performed using the steepest descent algorithm for 50,000 steps with a convergence criterion of 1000 kJ/mol/nm. The system was equilibrated in two phases: a 100 ps NVT ensemble at 300 K followed by a 100 ps NPT ensemble at 1 bar, during which position restraints were applied to heavy atoms. Temperature was maintained using the V-rescale thermostat (τ = 0.1 ps), and pressure was controlled using the Parrinello–Rahman barostat (τ = 2.0 ps). Long-range electrostatic interactions were treated using the Particle Mesh Ewald (PME) method with a real-space cutoff of 1.0 nm, a Fourier spacing of 0.125 nm, and a PME interpolation order of 4, while van der Waals interactions were truncated at 1.0 nm. Production simulations were carried out with a 2-fs time step. Production simulations were executed on a single NVIDIA GeForce RTX 5080 GPU using 1 thread-MPI process and 16 OpenMP threads, with GPU offloading of non-bonded interactions, PME, bonded interactions, and coordinate updates. All simulations were performed using GROMACS version 2025.4.

### 2.3 Trajectory analysis

Conformational stability and flexibility across all production trajectories was assessed through root-mean-square deviation (RMSD) and root-mean-square fluctuation (RMSF) analyses using standard GROMACS analysis modules. RMSD was calculated separately for histone protein backbone Cα atoms and DNA duplex heavy atoms, each fitted separately by least-squares alignment to the corresponding atoms of the energy-minimized reference structure. System equilibration and conformational convergence was evaluated through inspection of RMSD trajectories over time. Convergence was defined as the achievement of a stable RMSD plateau over the final 25% of the 200 ns production trajectory (150–200 ns). All 44 independent simulation runs attained stable plateau profiles within this window. For each of the 22 mutation pairs, plateau RMSD values were compared between WT and mutant simulations, and the difference (ΔRMSD = RMSD_mutant − RMSD_WT) was calculated for both protein and DNA components and correlated with MM-GBSA ΔΔG values using Spearman rank correlation.

Per-residue RMSF was calculated for DNA backbone heavy atoms, averaged over the full production trajectory after equilibration, to quantify position-specific conformational flexibility along the nucleosomal DNA. RMSF profiles were compared between wild-type and mutant systems for each mutation pair, and the per-residue difference (ΔRMSF = RMSF_mutant − RMSF_WT) was computed and visualized as a heatmap across all 22 mutation pairs to identify conserved patterns of mutation-induced flexibility change across the 146 bp nucleosomal DNA.

### 2.4 MM-GBSA binding free energy calculations

To calculate end-state free energy gmx_MMPBSA version 1.6.4 was used [18] [34]. The ff99SBildn force field (closest available to AMBER14SB in gmx_MMPBSA) was used to calculate the internal term (ΔE_int_), van der Waals (ΔE_VdW_), and electrostatic (ΔE_ele_) energies. Polar solvation free energies were calculated using the Onufriev–Bashford–Case Generalized Born model (GB-OBC2; igb = 5) [35], chosen for its optimal performance on protein-nucleic acid systems, with a physiological salt concentration of 0.15 M. The external dielectric was set to 78.5, and the internal dielectric was 2.0. The total net energy contributions were measured for each individual residue to allow precise determinations of the impact of each binding interaction. Interaction entropy was not tabulated due to numerical instability on large flexible systems.

Frames were analyzed at intervals of 50, after equilibration following start frame 1000. A relative mutation energy was calculated (ΔΔG calculation: ΔΔG = ΔG(mutant) − ΔG(WT)) for each paired simulation.

### 2.5 Statistical Analysis

All statistical calculations were executed in Python (v3.13.2) using the scipy.stats library. The net distribution of ΔΔG values across all 22 paired complexes was evaluated using a two-tailed Wilcoxon signed-rank test to assess whether the median ΔΔG differed significantly from zero. That is, whether oncogenic mutations produce a non-random net shift in nucleosome binding energy. The proportion of stabilizing versus destabilizing mutations was evaluated using a two-sided exact binomial test against a null expectation of equal probability (*p₀* = 0.5). Differences in binding energy shifts between the RAS oncogene family (KRAS, NRAS, HRAS) and kinase oncogenes (EGFR, BRAF, MAP2K1) were assessed using a two-sided Mann-Whitney U test.

Bivariate correlations between ΔΔG and structural parameters were computed using the Spearman rank-order correlation coefficient (*r_s_*). Statistical significance across all evaluations was established at *α* = 0.05.

## Results

### 3.1 The oncogenic mutation landscape

The 22 paired wild type/mutant simulations yielded ΔΔG values spanning approximately 147 kcal mol^-1^, from −59.4 kcal mol^-1^ (CDK4_24, strongly stabilizing) to +86.1 kcal mol^-1^ (EGFR_858, strongly destabilizing) (Figure 1). For an individual mutation to be statistically significant (*p* < 0.05), its mean value must be at least twice the size of its error bar, of which only one mutation present clears that threshold (EGFR 858). Even weak trends (*p* < 0.32) are only met by 7 of 22 mutations, while 15 of 22 mutations cross the zero baseline even at 1 × SEM. Consequently, rather than viewing these mutations as uniform disruptors, the data suggests a continuous thermodynamic spectrum best analyzed by distribution testing.

**Figure 1.**
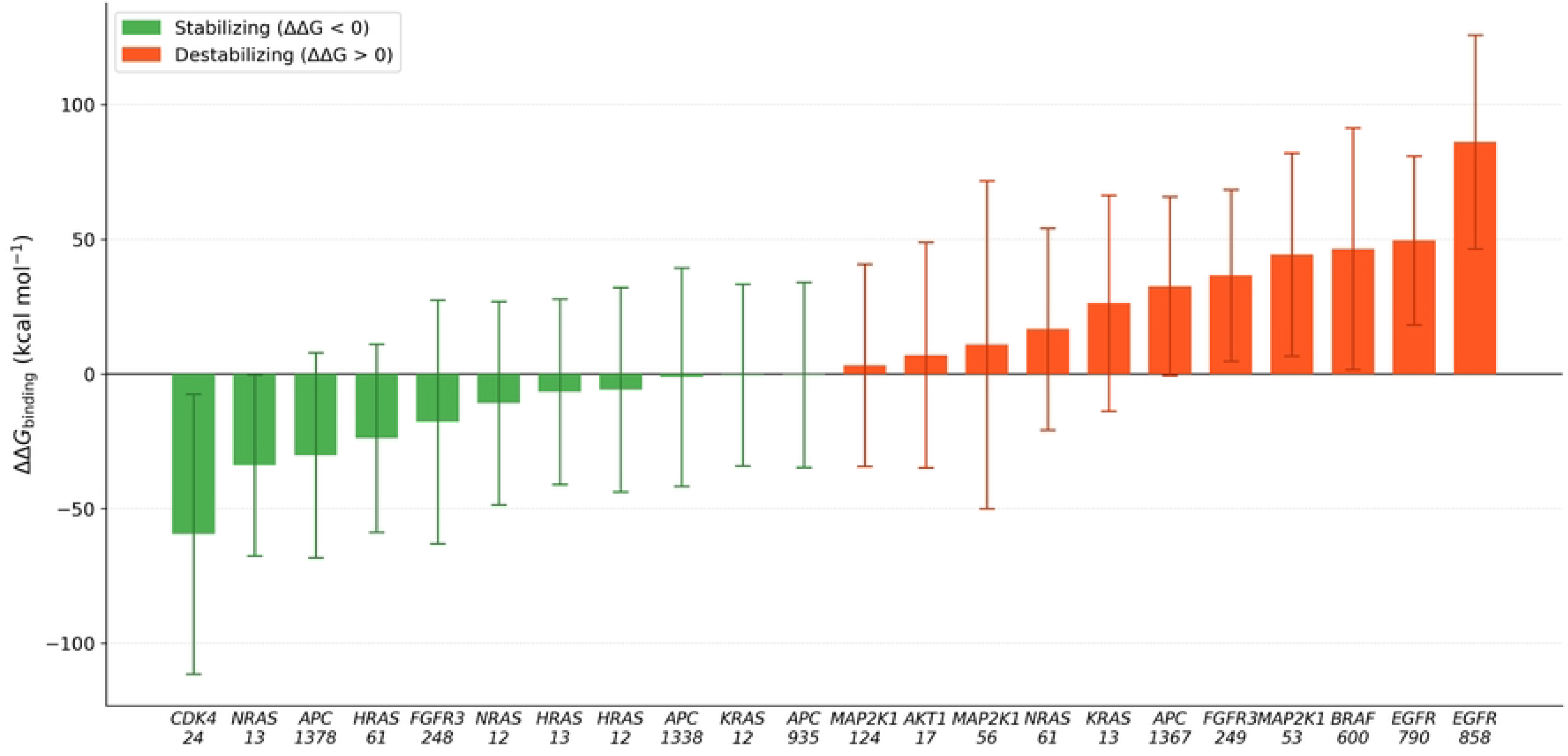
Impact of recurrent oncogenic somatic mutations on nucleosome core particle binding stability. Binding free energy shifts (ΔΔ*G*_binding_ = Δ*G*_mutant_ ― Δ*G*_wild type_) across 22 cancer hotspot mutations modeled in the human nucleosome core particle (PDB ID: 2CV5). Green bars indicate relative stabilization of the histone–DNA complex (ΔΔ*G* < 0 kcal mol^-1^), whereas red bars indicate relative destabilization (ΔΔ*G* > 0 kcal mol^-1^). Bars represent the mean binding free energy difference calculated via MM-GBSA across the stable plateau phase (150–200 ns; 381 frames), with error bars indicating standard error of the mean ( ± SEM). Mutations are ordered from most stabilizing to most destabilizing.

The distribution appears to be split by gene family, with RAS family mutations trending toward negative ΔΔG values and kinase mutations toward positive values. To test whether the overall distribution of ΔΔG values differed from zero, a two-tailed Wilcoxon signed-rank test was applied across all 22 mutations, yielding a non-significant result (*p* = 0.388). Similarly, the 11/11 split between stabilizing and destabilizing mutations did not differ from chance expectation (binomial test, *p* = 1.0). These non-significant results indicate that oncogenic mutations do not produce a uniform net shift in nucleosome binding energy across the full panel. Rather, the data suggest a bimodal distribution stratified by gene family, leading to the family-level analysis presented in Section 3.2.

### 3.2 Gene-family-dependent thermodynamic effects

Partitioning binding free energy shifts by oncogenic gene family revealed distinct, functional-class thermodynamic profiles (Table 1).

**Table 1.** Energetic impact of oncogenic mutations on nucleosome core binding stability grouped by gene family.

| Gene Family | N | Mean $\Delta\Delta G_{\text{binding}}$ (kcal<br>mol<sup>-1</sup>) | Direction |
| --- | --- | --- | --- |
| CDK4 | 1 | -59.4 | All stabilizing |
| HRAS | 3 | -12.1 | All stabilizing |
| NRAS | 3 | -9.4 | Predominantly stabilizing |
| APC | 4 | +0.3 | Neutral/mixed |
| AKT1 | 1 | +7.0 | Destabilizing |
| FGFR3 | 2 | +9.4 | Mixed |
| KRAS | 2 | +12.9 | Mixed trending destabilizing |
| MAP2K1 | 3 | +19.5 | All destabilizing |
| BRAF | 1 | +46.5 | Destabilizing |
| EGFR | 2 | +67.8 | All destabilizing |

The RAS family (KRAS, NRAS, HRAS) in this dataset showed a trend toward nucleosome stabilization, with a mean ΔΔG = −4.82 kcal mol^-1^. On the other hand, kinase domain mutations (EGFR, BRAF, MAP2K1) trended toward destabilization, with a mean ΔΔG = +40.12 kcal mol^-1^. Using a Mann-Whitney U test gives *p* = 0.008, suggesting a relationship between gene family and thermodynamic direction of effect in this dataset.

While all NRAS and HRAS mutations were consistently stabilizing, KRAS exhibited divergence. The KRAS_12 mutation was stabilizing (ΔΔ*G* = ―0.4 kcal mol^-1^), and KRAS_13 was destabilizing (ΔΔ*G* = +26.3 kcal mol^-1^). While all 3 of these GTPases share similar catalytic core domains, their biological functions are distinct. They each have different preferences for downstream effectors. The sequence context of KRAS leads to a nonuniform nucleosome energy landscape, demonstrating that even within a specific family, localized sequences alter the thermodynamic effects. This within-family divergence suggests that even single nucleotide differences in sequence context can override family-level thermodynamic trends, and highlights KRAS as a complex case that warrants further investigation.

### 3.3 VdW-driven mechanism and interface localization

MM-GBSA energy decomposition revealed that the raw electrostatic and polar solvation components were large in magnitude, but opposing in sign across all 22 mutation pairs. The raw electrostatic and polar solvation components individually exceeded several hundred kcal mol^-1^ in absolute value, though these are not shown directly as their net sum is the meaningful quantity (Supplementary Table S1). This near-complete cancellation is a well-established physical consequence of the electrically charged nature of DNA-protein interfaces: improvements in electrostatic complementarity between the negatively charged DNA backbone and positively charged histone residues are offset by the energetic cost of displacing structured water from the interface (desolvation penalty). The net electrostatic contribution (the sum of the electrostatic and polar solvation components) was therefore substantially smaller in magnitude than either component in isolation.

After accounting for this cancellation, the net energetic drivers of ΔΔG_binding varied across the mutations. Across the full survey, VdW changes were the primary driver in 12 of 22 mutations, while residual net electrostatics dominated in the remaining 10 (Supplementary Table S1). Most significantly, the results on the extreme ends of the spectrum were dominated by VdW forces, while the mutations with moderate ΔΔG values showed electrostatics forces more comparable in size to VdW forces (Figure 2). Non-polar solvation contributions were consistently small (range: −5.5 to +5.7 kcal mol^-1^) and did not drive the overall ΔΔG in any mutation pair. This pattern is most clearly demonstrated by the mutations producing the largest thermodynamic effects: CDK4_24 (ΔΔG = −59.4 kcal mol^-1^) and EGFR_858 (ΔΔG = +86.1 kcal mol^-1^), for which the van der Waals (VdW) component was the dominant contributor (accounting for −57.1 and +60.5 kcal mol^-1^ respectively), with the net electrostatic contribution being comparatively modest (+3.1 and +20.0 kcal mol^-1^) (Supplementary Figures S2 and S3).

**Figure 2.**
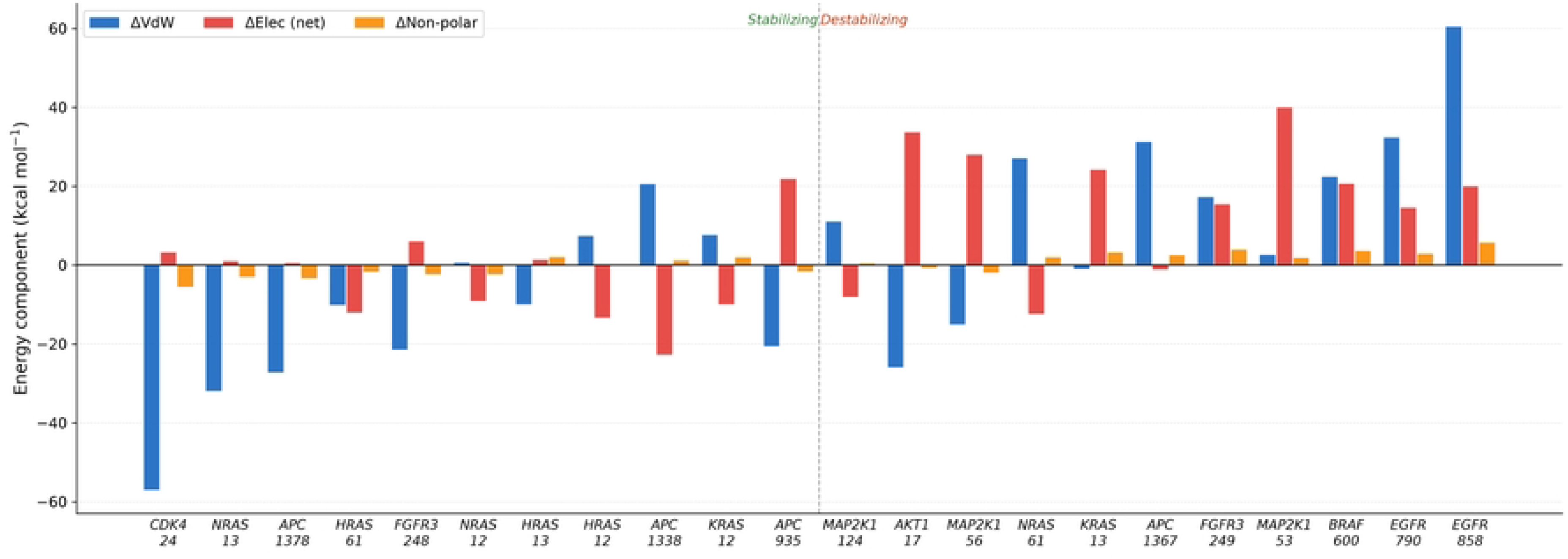
Energetic decomposition of binding free energy shifts across 22 oncogenic mutations. Decomposition of MM-GBSA relative binding free energies (ΔΔ*G*_binding_) into van der Waals (ΔΔ*E*_vdW_, blue), net electrostatic (ΔΔ *E*_elec(net)_ = ΔΔ*E*_elec_ + ΔΔ*G*_GB_, red), and non-polar solvation (ΔΔ*G*_non-polar_, orange) contributions. Mutations are ordered from most stabilizing (left) to most destabilizing (right) following Supplementary Table S1. High-magnitude stabilization and destabilization extremes are predominantly governed by VdW interactions, whereas electrostatic contributions dominate intermediate, ambivalent variants.

### 3.4 Interface localization

Per-residue MM-GBSA was run exclusively to measure interactions within 6 Ångströms of the binding partner, thereby only measuring interactions at the interface between DNA and the histones. The interface capture percentage (Supplementary Table S1) is how much of the total ΔΔG is explained by just the interface residues, where 100% means that the entire stability is driven by interface interactions. If the value is much lower than 100% it would mean a substantial amount of the energy change is happening far from the interface, as the mutation is acting through long range effects.

Among the 18 mutations with |ΔΔG| > 5 kcal mol^-1^, where the capture fraction is numerically reliable, the mean interface capture was 104.3% (median 98.0%, range 61.0–164.0%), indicating that the thermodynamic effects of these mutations are overwhelmingly localized to within 6 Å of the DNA-histone interface. The four near-neutral mutations (|ΔΔG| < 5 kcal mol^-1^) were excluded from this calculation because division by a near-zero denominator produces numerically unstable capture fractions (Figure 3). This result indicates that these mutations change how DNA and histones bind to each other by altering specific local contacts at the surface where they touch, and not by causing global structural rearrangements throughout the whole nucleosome. This correlates with the results in Section 3.3 that the mechanistic component that explains the stability changes is VdW forces.

**Figure 3.**
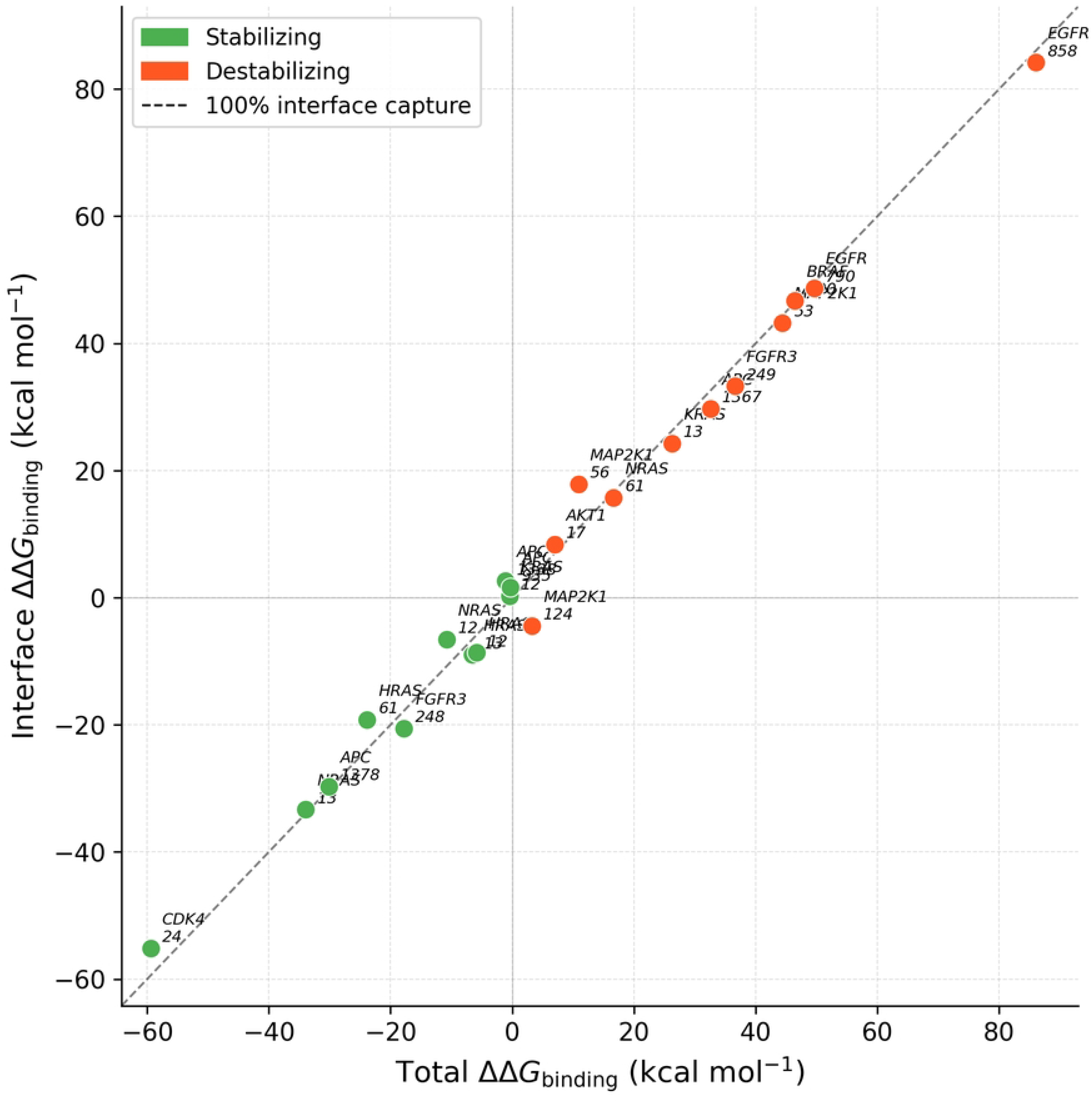
Interface localization of mutation-induced binding free energy changes. Total ΔΔG_binding (x-axis) plotted against the sum of per-residue ΔΔG contributions from residues within 6 Å of the DNA–histone interface (y-axis) for all 22 mutation pairs. The dashed diagonal represents 100% interface capture. Points falling on or near the diagonal indicate that the thermodynamic effect of the mutation is fully accounted for by interface residues alone.

### 3.5 RMSD and RMSF patterns

To confirm the reliability of the MM-GBSA free energy values, the conformational stabilities of all 44 simulations were measured through backbone RMSD analysis. All 22 mutation pairs reached stability within the first quarter of their respective simulations (Supplementary Figure S4, Supplementary Table S2). All 22 pairs maintained the stable plateau during the 150-200 ns of the simulation (the portion used for MM-GBSA). Protein backbone RMSD plateaued at a mean of 1.39 Å (range: 1.30–1.49 Å) for wild-type simulations and 1.43 Å (range: 1.30–1.69 Å) across mutant simulations. This mean difference of only +0.04 Å confirms that the histone octamer structure was well-preserved throughout all simulations regardless of the mutation.

DNA RMSD plateaued at a mean of 2.98 Å (range: 2.52–4.04 Å) for wild-type simulations and 2.90 Å (range: 2.49–3.51 Å) for mutant simulations, consistent with the greater conformational flexibility expected of a 146 bp nucleosomal DNA compared to the compact histone. The small range of protein plateau RMSD values across all 22 wild-type simulations (1.30–1.49 Å) demonstrates that the nucleosome core particle structure is inherently stable for the simulation conditions utilized and that across simulation variability does not introduce systematic structural drift.

Per-residue RMSF showed different DNA flexibility dependent on the location within the nucleosome, visualized as a heatmap of ΔRMSF (mutant minus wild-type) (Figure 4).

**Figure 4.**
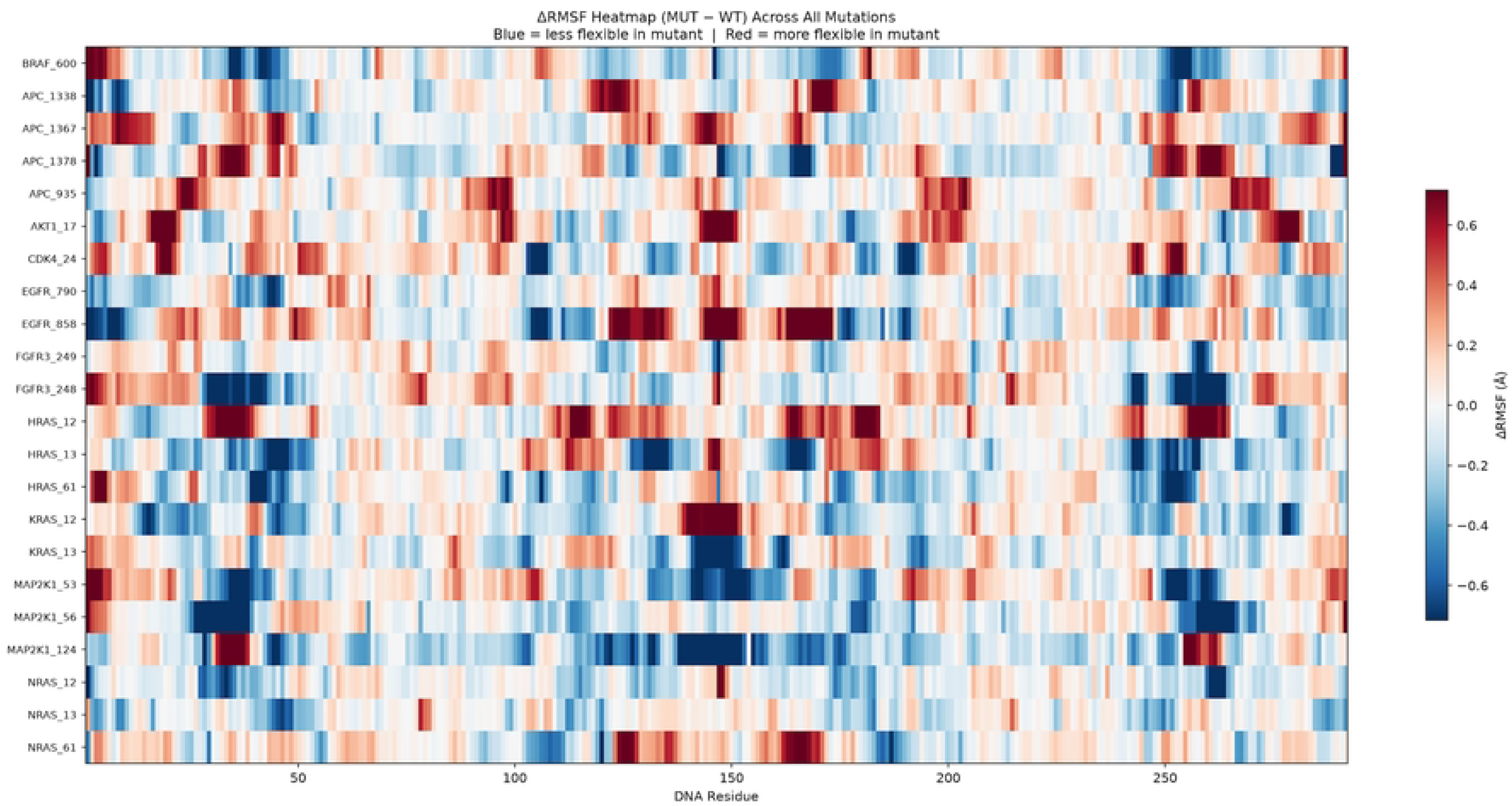
Heatmap of per-residue DNA RMSF differences (ΔRMSF = RMSF_mutant − RMSF_WT) across all 22 oncogenic mutation pairs. Each row represents one mutation pair, and each column represents one DNA residue position along the 146 bp nucleosomal DNA. Red indicates positions where the mutant DNA is more flexible than wild type; blue indicates reduced flexibility. Mutations are ordered by ΔΔG_binding from most stabilizing (top) to most destabilizing (bottom).

There was little reduction in mean DNA flexibility (mean ΔRMSF = −0.021 Å), but individual mutations produced substantial localized changes. AKT1_17 had an increase of +2.61 Å (largest increase) while MAP2K1_124 had a decrease of −4.56 Å (largest decrease). AKT1_17 also had the greatest DNA RMSD increase of any mutation (ΔRMSD = +0.71 Å), suggesting broader conformational disruption relative to other destabilizing variants. The three MAP2K1 mutations showed a consistent pattern of DNA rigidification (mean ΔRMSF: −0.29, −0.14, and −0.07 Å for MAP2K1_124, _56, and _53 respectively), aligning with their consistent destabilizing thermodynamic profile (Section 3.2). Notably, the two EGFR mutations showed divergent structural behavior: EGFR_790 produced net DNA rigidification (ΔRMSD = −0.79 Å) while EGFR_858 showed increased flexibility (ΔRMSD = +0.25 Å). This suggests mechanistically distinct structural pathways to energetic destabilization within the same gene family (Supplementary Figure S5).

Spearman correlation analysis revealed no significant association between ΔRMSD and ΔΔG_binding for protein (ρ = 0.228, *p* = 0.309), DNA (ρ = 0.001, *p* = 0.998), or mean ΔRMSF (ρ = 0.051, *p* = 0.820). This absence of correlation is consistent with the interface localization finding in Section 3.4 that thermodynamic effects arise from precise local contact changes at the DNA–histone interface rather than global conformational rearrangements.

## Discussion

### 4.1 Gene-family-dependent chromatin effects and oncogenic mechanisms

Hotspot recurrence in cancer reflects both positive selection for functional advantage and intrinsic mutational susceptibility [5, 36]. In the dataset examined, the RAS family gene mutations trended toward stabilizing the DNA-histone interactions. A more stable nucleosome at the RAS genes could inhibit the access to repair enzymes that can correct oncogenic mutations. Nucleosome-bound DNA is repaired less efficiently because repair enzymes have reduced access, and more tightly bound mutant DNA may therefore be harder to repair. This creates a cycle in which the mutant DNA then becomes more challenging to repair by very nature of the mutation. This mechanism is consistent with the research by Schuster-Böckler and Lehner [37] and Pich et al [15], who demonstrated that mutation rates are elevated at certain nucleosome positions. Additionally, tightly compacted chromatin is associated with late replication timing, and late replication regions demonstrate elevated mutation rates [38]. Together, these mechanisms may contribute to increased levels of mutations at certain RAS hotspots. However, kinase domain mutations in the dataset consistently trended towards destabilizing the nucleosome complex. Unlike RAS, which leads to oncogenesis through constitutive protein activation, kinase driven cancers such as EGFR driven tumors involve increased expression of the oncogenic protein through transcriptional upregulation. Therefore, greater chromatin accessibility may support continued oncogene protein expression. This is consistent with the established role of chromatin remodeling and enhancer accessibility in EGFR-driven cancers, where open chromatin and active enhancer regions contribute to EGFR-associated transcriptional activity [39]. The opposing thermodynamic effects observed for the RAS and kinase families may therefore reflect fundamentally different dependencies on the chromatin environment. RAS stabilization may reinforce mutation persistence by preventing DNA repair, while kinase destabilization facilitates transcriptional upregulation of the oncogenic kinase. Whether these thermodynamic patterns have functional consequences at the actual genomic loci of these genes in vivo remains to be determined experimentally. The present findings reflect the biophysical consequences of the nucleotide substitutions characteristic of each oncogene family when placed in a standardized context. Critically, the finding that thermodynamic effects are overwhelmingly localized to within 6 Å of the DNA-histone interface across 18 of 22 mutations provides mechanistic specificity to these observations. These mutations act through precise local contact changes at defined positions, not through structural rearrangements. This localization makes the predicted effects directly testable through site-specific mutational analysis of interface residues

MM-GBSA has inherent limitations in its ability to calculate individual ΔΔG values. However, this does not detract from its ability to accurately order and rank relative ΔΔG values, as supported by the Mann-Whitney U test. The MD simulations performed used a single nucleosome structure and do not consider the overall chromosomal structure present in cells.

Additionally, the DNA locations on the nucleosome were determined with a predictive model (NuPoP). Future studies using locus-resolved MNase-seq or histone ChIP-seq datasets to validate nucleosome positions would strengthen the results. This study does not model the dynamic processes that occur *in vivo*; rather, the 200 ns simulation captures local conformation dynamics. The predictions from this study are directly testable using *in vitro* nucleosome stability assays.

While it has long been understood that protein-level functional advantage selects for recurring oncogenic mutations [36], our data suggest that these substitutions also alter the local thermodynamic stability of the nucleosome complex in a gene-family dependent manner. It is further possible that there are selective pressures inherent in the thermodynamic stability of the nucleosome that also select for oncogenic mutations. Local thermodynamic effects at the nucleosome level may therefore not be merely an accessory byproduct to a mutation, but potentially a contributing pressure that influences mutations. The observation that somatic mutations may alter nucleosomal thermodynamics in a specific manner highlight chromatin stability as an underappreciated biophysical component in tumor evolution, which begs for further exploration.

## Acknowledgments

The authors thankfully acknowledge Provost Selma Botman of Yeshiva University for graciously providing funding for this research.

## Supporting Information

**Supplementary Figure S1. Crystal structure of the nucleosome core particle (PDB ID: 2CV5).** The 146 bp nucleosomal DNA double helix (strands colored orange and green) is shown wrapped around the histone octamer core (subunits shown in cartoon representation). This crystallographic conformation served as the baseline structural template for all molecular dynamics simulations.

**Supplementary Figure S2. Per-residue MM-GBSA energy component decomposition for CDK4_24 (ΔΔG = −59.4 kcal mol^-1^).** Bars show the ΔΔG contribution from each significant interface residue, decomposed into VdW (blue), electrostatic (red), polar solvation (green), and non-polar solvation (orange) components. Negative values indicate residues that contribute to stabilization in the mutant relative to wild type. The dominance of the polar solvation component (green) reflects the large electrostatic–solvation cancellation described in Section 3.3. The net thermodynamic effect after cancellation is primarily VdW-driven.

**Supplementary Figure S3. Per-residue MM-GBSA energy component decomposition for EGFR_858 (ΔΔG = +86.1 kcal mol^-1^).** Bars show the ΔΔG contribution from each significant interface residue, decomposed into VdW (blue), electrostatic (red), polar solvation (green), and non-polar solvation (orange) components. Positive values indicate residues that contribute to destabilization in the mutant relative to wild type. As with CDK4_24, the large polar solvation bars reflect the electrostatic–solvation cancellation inherent to charged DNA-protein interfaces. The net destabilizing effect is primarily VdW-driven at the thermodynamic extremes.

**Supplementary Figure S4. Structural stability and backbone RMSD equilibration across 200 ns trajectories.** Time-series backbone root-mean-square deviation (RMSD) profiles for wild-type (blue) and mutant (red) nucleosome complexes. The shaded gray window denotes the final 25% production plateau analyzed for binding free energy calculations. **(A)** CDK4_24, representative stabilizing mutation, histone octamer core (1.48 ± 0.05 Å WT vs. 1.48 ± 0.07 Å Mutant). **(B)** CDK4_24 nucleosomal DNA (2.93 ± 0.18 Å WT vs. 3.25 ± 0.16 Å Mutant). **(C)** EGFR_858, representative destabilizing mutation, histone octamer core (1.38 ± 0.06 Å WT vs. 1.41 ± 0.08 Å Mutant). **(D)** EGFR_858 nucleosomal DNA (3.01 ± 0.18 Å WT vs. 3.26 ± 0.25 Å Mutant).

**Supplementary Figure S5. Per-residue DNA RMSF profiles for EGFR_790 and EGFR_858, depicting divergent structural mechanisms within the same gene family.** (A) EGFR_790 (ΔΔG = +49.6 kcal mol^-1^). Top panel: RMSF values for wild-type (blue) and mutant (red) across all nucleosomal DNA residues. Bottom panel: per-residue ΔRMSF (mutant − wild-type) where green bars indicate positions where the mutant DNA is less flexible than wild-type (below −1σ); and orange bars indicate increased flexibility (above +1σ). EGFR_790 shows predominantly negative ΔRMSF values consistent with net DNA rigidification (ΔRMSD = −0.79 Å; 58 residues less flexible). (B) EGFR_858 (ΔΔG = +86.1 kcal mol^-1^). Despite producing a larger total destabilization, EGFR_858 shows a more balanced RMSF profile with localized regions of increased DNA flexibility (ΔRMSD = +0.25 Å; 43 residues more flexible). The opposing ΔRMSF signatures of EGFR_790 and EGFR_858 indicate that these two hotspot mutations destabilize nucleosome binding through structurally distinct mechanisms despite qualitatively similar thermodynamic outcomes.

**Supplementary Table S1.** MM-GBSA binding free energy components for all 22 oncogenic mutation pairs.

**Supplementary Table S2. Plateau RMSD and mean RMSF statistics for all 22 mutation pairs.**

